# Critical analysis of the mosquito repellency evoked by the 10-34 kHz recorded animal sounds: The case of the African female *A. gambiae s.s*

**DOI:** 10.1101/841387

**Authors:** Philip Amuyunzu Mang’are, Francis Ndiritu Gichuki, Samwel Rotich, Jacqueline K. Makatiani, Bernard Rapando Wakhu

## Abstract

Animals sounds have been mimicked in electronic mosquito repellents (EMRs) and exploited as a tool in the control of malaria by targeting the vector, the female *Anopheles gambiae s.s.* The claimed mosquito repellency of 30.3 % due to Anti-Pic^®^, an electronic mosquito repellent, had failed to be confirmed in subsequent studies. However, studies on mosquito startle based on initial behavioural activities without an attractant yielded 34.12 % repellency elicited by the 10-34 kHz recorded sound of *O. tormota*. Other malaria intervention measures involving the use of chemicals have been impeded by the pathogen and vector resistance hence slowing down the rate of decline of malaria morbidity and mortality. The research thus focused on the analytical study of the African female *A. gambiae s.s* repellency evoked by the 10-34 kHz recorded animal sound of male mosquito, *Anopheles gambiae* and *Delphinapterus leucas*. Landing rates and behavioural startle responses of the mated female *A. gambiae* on food attractant evoked by the individual sound of the male mosquito, *A. gambiae*, *O. tormota* and *D. leucas* were determined and analysed. The male and female *A. gambiae* were bred and reared under controlled laboratory conditions of 60-80 % humidity, 25±2 °C temperature with equal light-darkness hour cycle in KEMRI, entomology laboratories. Isolation of the male and female mosquitoes from a swarm was based on physical features and affinity to blood meal. The sounds of *O. tormota* and *D. leucas* were acquired and the sound of the male *A. gambiae* were recorded from the Kenya Medical Research Institute (KEMRI) entomology laboratory, Kisumu. The sounds were filtered into 10-34 kHz frequency band and analysed using Avisoft-SAS LAB Pro version 5.2 and Raven Pro 1.5 software. The sound of *O. tormota* was also studied. A fighto-Y glass cage well designed into control, neutral and treatment chambers was used in the study. Both control and treatment chambers were connected to blood meal maintained at 38.60°C. The treatment cage was also connected to the source of sound and a swarm of 50 female mosquitoes into the neutral cage and observed for 1,200 s. The sounds of the *A. gambiae, O. tormota* and *D. leucas* yielded 2.10, 2.20 and 3.00 landings/minute respectively associated with adverse behaviour. The protection index (PI) anchored on the number of mosquitoes that landed, probed and fed on the blood meal in the treatment and neutral cage for the sounds of the *A. gambiae, O. tormota* and *D. leucas* was 42.73 %, 40.24 % and 10.64 % respectively. The sound of the *A. gambiae* was characterised by steady and minimally dipped pulsate acoustic power with wide bandwidth. The protection index achieved by the sound of the male *A. gambiae* did not differ significantly from the sound of *O. tormota* (0.1740 > 0.05), though differed significantly from the sound emitted from the Anti-Pic^®^ EMR (p = 5.3440 x 10^−5^).

**The author summary:** Philip Amuyunzu Mang’are is a PhD. Physics student in Egerton University. He has authored many papers and books. He is currently a Lecturer of Physics (Electronics), Masinde Muliro University of Science and Technology. He is a member of the Biophysical Society and the current President of Biophysical society (Kenya). Prof. Ndiritu F. Gichuki, is a Professor of Physics Egerton University. Currently he is the Registrar Academic Affairs in Chuka University. His vast experience has seen him supervise many postgraduate students who have taken key positions in the society. Prof. Samwel Rotich is a Profesor of Physics in Moi University specialising in Electronics. He has a wide experience in Physics and Biophysics. He is a registered member of the Biophysical Society and the Patron of Biophysical Society Kenya Chapter. He has published many papers and supervised many postgraduate students. Dr. Makatiani Kubochi is a Lecturer in Moi University with vast experience in entomology. She has published many papers and supervised many postgraduate students. Dr. Rapando Bernard Wakhu is a renown theoretical Physicist with experience in acoustics and Fourier analysis based in Masinde Muliro University of Science and Technology. He has supervised many postgraduate students and published many papers.

## 1. Introduction

### 1.1 Highlights of the Worldwide Malaria Situation and Interventions

Adherence to recommended interventions makes malaria a preventable and treatable disease (WHO, 2011). Malaria interventional approaches targeting the vector and which involve the use of insecticide-treated nets (ITNs), indoor residual spraying (IRS) and in some specific settings, larval control are a critical components of the multipronged attack on malaria (Godfray, 2013). The chemoprevention is used for the most vulnerable populations, particularly pregnant women and infants. Confirmation of malaria diagnosis through microscopy or rapid diagnostic tests (RDTs) for every suspected case and timely treatment with appropriate antimalarial medicines is key in malaria control (CDC, 2010; Enayati and Hemingway, 2010; WHO, 2011; Godfray, 2013; WHO, 2013; WHO, 2015; Gething *et al*., 2016).

Protection by ITNs and IRS has demonstrated greater impact in reducing malaria (WHO, 2013). Hwever, strains of *Anopheles* mosquitoes developed resistance to DDT, pyrethroid and other insecticides, and the environmental impact of DDT was recognized. Also, the *Plasmodium* parasites became resistant to chloroquine, the mainstay of antimalarial drug treatment in humans (WHO, 2006; Enayati and Hemingway, 2010). Sound has over the years been used to scare off pest species, with its humble origins of loud claps and yells in ancient agricultural fields, and now ultrasound producing electronic repellents (EMR) are used in combating mosquitoes (Antonelli *et al.*, 2007; Enayati *et al*., 2010; Simeon *et al*., 2013; Aflitto and DeGomez, 2014). Evaluation experiments with mosquito-repelling devices which included the Anti-Pic^®^, Mosquito Repeller^®^ DX-600 and Bye-Bye Mosquito^®^ done by exposing human hands to *Aedes albopictus* (Skuse) adults showed insignificant success in repellency, failing to confirm the 30.3 % repellency due to Anti-Pic^®^ initially determined (Andrade and Bueno, 2001). The electronic mosquito-repelling devices studied ranged from 2kHz to 60 kHz in frequency with harmonic peaks from 4 kHz to 68 kHz (Rutledge *et al*., 1985; Combemale *et al*., 1992). Also, experiments with functioning electronic mosquito repellents (EMR) mimicking calls from bats and male *Anopheles gambiae* in the frequency range of 125 Hz to 74.6 kHz showed that 12 out of 15 field experiments yielded higher landing rate on the human bare body parts than the control experiments (Andrade and Bueno, 2001; Center For the Advancement Of Health, 2007; Enayati *et al*, 2010). The EMRs are used indoors and outdoors and are purported to repel mosquitoes within a range of 2.5 m (CAH, 2007). Recent researches with natural and synthetic sounds have shown startle response in mosquitoes, thus sound was an effective additional tool in the control of mosquitoes, the malaria vector (Mohankumar, 2010; Mang’are *et al*., 2012). Evasive responses characterized by 58.5° antenna erection, physical injury, fatigue and falls; attributed to pronounced stress on the nervous system and fear of predation have been reported for both 10-34 kHz and 35-60 kHz sounds of *Odorrana tormota* based on behaviural activity. An average percentage startle of response of 34.12 % and 46 % was observed in mosquitoes in the 10-34 kHz and 35-60 kHz sounds of *O. tormota* respectively (Mang’are *et al*., 2012). Researches based on mosquito landing, probing and feeding determined the percent Protection Index (PI) or Repellency using the equation 1.1 (Rutledge *et al*., 1985; Combemale *et al*., 1992):

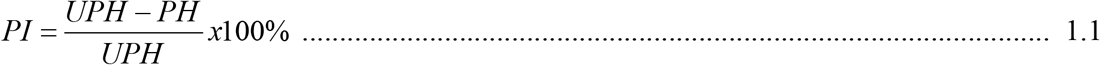

where ***PH*** is the number of mosquito bites (or initiated bites) on the supposedly protected blood meal (treatment), and ***UPH*** is the same measure for supposedly unprotected blood meal (control).

Other studies have established that mosquitoes detect ultrasound in the range of 38 - 44 kHz, regardless of the source, initiating avoidance response since it creates stress on their nervous system, jams mosquitoes’ own ultrasound frequency besides immobilizing them (Mohankumar, 2010). The startle response in mosquitoes evoked by the natural sound of the male, *Delphinapterus leucas and Tursiops truncatus* had not been reported. The sounds of *A. gambiae* and *O. tormota* had been studied hence the basis for this study.

### 1.2 Mosquito biology and Audition

The egg, larva and pupa stages in the lifecycle of the *A. gambiae* are aquatic and last 5-14 days, depending on the species and the ambient temperature (CDC, 2010). Both male and female adult mosquitoes feed on plant nectar, but the female feed on vertebrates’ blood, for nutrients required for egg maturation (Foster and Walker, 2009). Warm-blooded hosts provide mosquitoes with thermal contrast that facilitates the localization of a suitable blood meal (McMeniman *et al.,* 2014; EarthSky in Earth, 2015). An analysis of how the mosquito actually bites, probes for the blood vessels and finally sucks bloods showed that the mean time taken before the mosquito starts probing after landing was 6.5 seconds, the mean probing time was 142 seconds, mean feeding time was 240 seconds (feeding times were between 150 and 329 seconds) giving a total of 389 seconds (6.5 minutes) (Choumet *et al.,* 2012). Female Anopheles mosquitoes lay eggs on the surface of the water at night and under favorable conditions, hatching occurs within one or two days and develop within the aquatic habitat. The *A. gambiae* larvae develop in permanent man-made structures and natural pools (Kweka *et al.,* 2012). The adult mosquitoes have slender bodies consisting of the head, thorax and abdomen; the head specialized for acquiring sensory information and for feeding. The mosquito antennae detects host and breeding sites odors (CDC, 2010). The head also has an elongated, forward-projecting proboscis used for feeding, and two sensory palps. The adult stages of many mosquito species are feeders of blood, which has given some disease causing organisms a reliable mode of transmission to animal hosts. Adult males and females *Anopheles* rest with their abdomens sticking up in the air. It is during the adult stage that the female *Anopheles* mosquito acts as malaria vector. The adult females can live up to a month or more in captivity but they don’t live more than 1-2 weeks in nature (Antonelli *et al.,* 2007). The mosquito has a pair of large, wraparound eyes, and a pair of long, hairy antennae; its ears projecting from the front of its face (Hoy, 2006). The antenna detects the particle velocity component of a sound field, which is restricted to the immediate vicinity of the sound source in acoustic near field. Male mosquitoes require about 24 hours before their terminalia get rotated and their fibrillae mature enough to become erect and detect females whereas the female mosquitoes need 48-72 hours before they become receptive to males prior to blood feeding in the wild (Clements, 1992). Anopheles males can mate several times, but females become refractory to re-insemination and re-mating is rare (Mohankumar, 2010). Mating in *anopheline* mosquitoes occur during the early evening, primarily in swarms (Diabaté *et al.,* 2011). The swarming males use their erect antennal fibrillae to detect a nearby female mosquito’s wing beat frequencies (Clements, 1992). The case study on Toxorhynchites showed that the males harmonized their wing beat with females as they neared, for species recognition, before mating commenced (Gibson and Russell, 2006). Flying mated female mosquitoes produce familiar whining sound when searching for proteins (Maweu *et al.,* 2009). This sound is very critical in this study as an audio switching system. Ultrasound generated artificially or naturally is detected by mosquitoes evoking evasive response (Mohankumar, 2010).

### 1.2 Statement ofthe Problem

Africa and the world as a whole suffers both economic and health burden due to malaria. Interventions which includes the use of chemicals targeting malaria pathogens and vectors have led to a decline in malaria mortality and morbidity though at a lower rate due to buildup of resistance. Also the use of electronic mosquito repellents mimicking the sounds of bats or male mosquitoes in the control of mosquitoes has been a debatable issues since the 30.3 % claimed repellency failed to be confirmed. However, studies on mosquito startle involving the use of recorded sound of *O. tormota* have been carried out yielding an average startle of 34.12 % and 46 % in mosquitoes in the 10-34 kHz and 35-60 kHz frequency bands. These startle values were determined from initial behavioural responses without an attractant. Therefore there was need to use an improved bioassay setup to determine and analyse the landing rates and behavioural startle responses of the mated female *A. gambiae* on food attractant evoked by the individual sound of the male mosquito, *A. gambiae, O. tormota. truncatus* and *D. leucas.* The data collected was based on the number of mosquitoes that approached the attractant; landed; landed and probed; or landed, probed, and bit or fed the repellent-treated blood meal in the treatment bioassay cages and the untreated meal in the Control chamber. The protection index PI against mosquitoes due to the animal sounds was determined and compared. The results determined confirmed the feasibility of using recorded animal sound as an additional malaria intervention measure.

### 1.3. Objectives

#### 1.3.1. General Objective

Investigate the startle response of the African female *Anopheles gambiae* by natural sound of male mosquito, *Anopheles gambiae, Tursiops truncatus, Odorrana tormota* and *Delphinapterus leucas*

#### 1.3.2. Specific Objectives

i. Determine the landing rates and behavioural startle responses of the mated female *A. gambiae* on food attractant evoked by the individual sound of the male mosquito, *A. gambiae*, *O. tormota, T. truncatus* and *D. leucas*.
ii. Analyse the landing rates and behavioural startle responses of the mated female *A. gambiae* on food attractant evoked by the individual sound of the male mosquito, *A. gambiae*, *O. tormota, T. truncatus* and *D. leucas*.

## 2. Materials and Methods

### 2.1. Sound of the *O. tormota,* male *A. gambiae, T. truncatus and D. leucas*

The male and female mosquitoes, *A. gambiae* s. s were bred and reared in Kenya Medical Research Institute (KEMRI) Entomology laboratories under controlled laboratory conditions. The pupae of *A. gambiae* s. s were reared in vials quarter filled with water and covered with a net at 60-80 % humidity, 25±2 °C temperature and equal light-darkness hour cycle at KEMRI/CDC, entomology laboratories. The male and female mosquitoes were discriminated based on physical features and affinity to blood meal. The recording of mosquito sounds and the bioassay studies was also conducted in KEMRI/CDC entomology. The sounds of the *O. tormota* recorded from Huangshan Hot Springs in Anhui Province, China were acquired from Prof. Albert Feng whereas the sounds from the *D. leucas* and *T. truncatus* were acquired from Prof. Herve Glotin of Institut Universitaire de France. The recorded sounds of the *O. tormota, male A. gambiae, T. truncatus* and *D. leucas* were used in the study individually and in combination. However, the sound of *T. truncatus* could not be involved in the bioassay study due to its inferior acoustic parameters. Hence the sounds of the *male A. gambiae* and *D. leucas* whose frequencies, amplitudes, bandwidths parameters and other spectral features were superior or matched closely with the sound of *O. tormota* were studied. The sound of *O. tormota* was also involved in the bioassay due to the improved bioassay setup and observations based on landing and probing of blood meal. The individual sounds of *O. tormota, male A. gambiae* and *D. leucas* were filtered using the Avisoft-SAS LAB Pro version 5.2 band-pass filters into 10-34 kHz, 35-60 kHz and 61-90 kHz frequency bands. The standard Fighto-Y bioassay glass cage fitted with a mosquito netting on the three cross-section areas, shown in Figure 1 was used in the study. Cotton wool was used to seal the entry/ exit hole on the top part of the glass cage.

**Figure 1:**
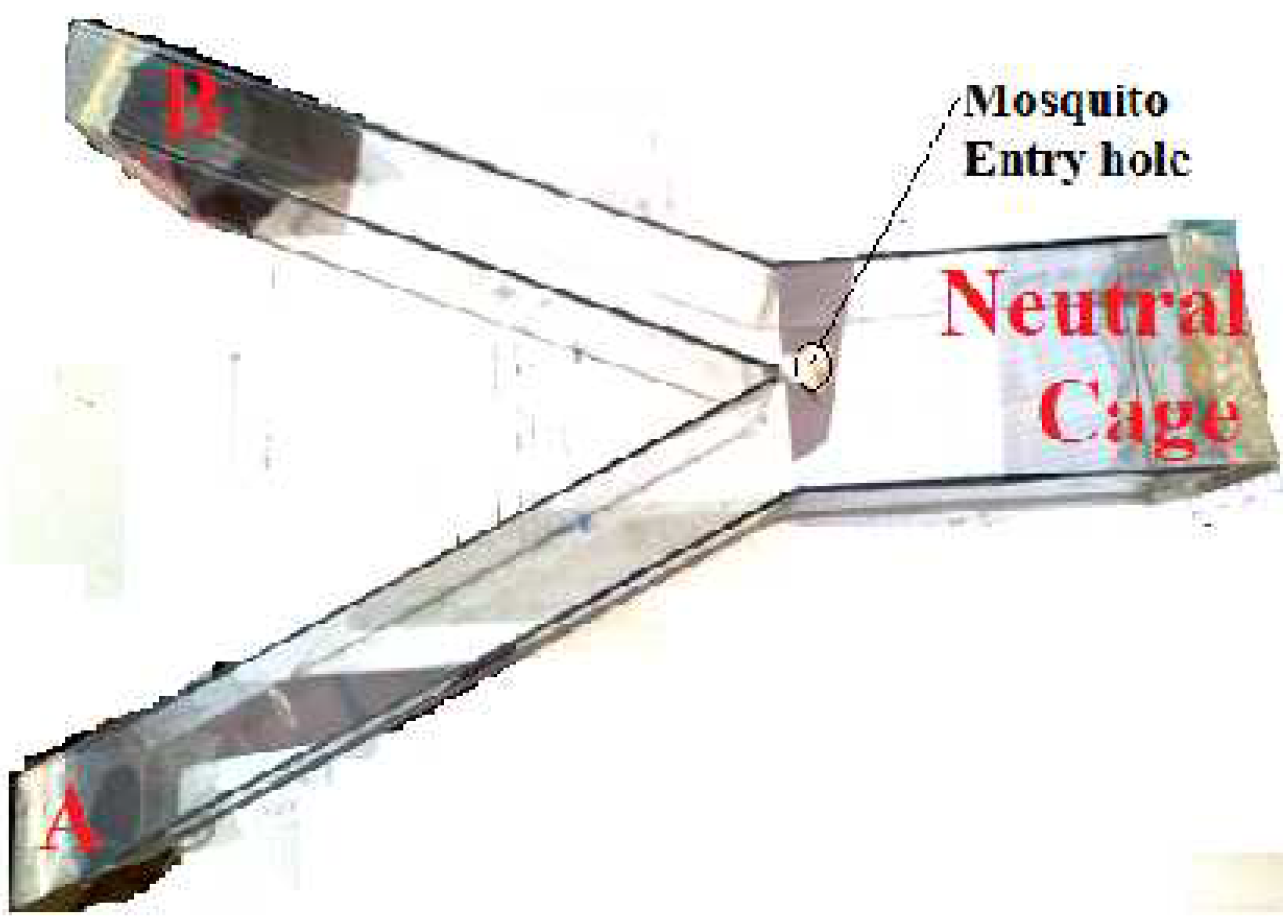
The Fighto-Y glass cage

The cage was divided into three sections, A, B and the Neutral cage. Both ends of cage A and B were open but covered with a mosquito netting. Similarly, the neutral cage had an open end (5 inch x 5 inch) was covered by a mosquito netting. Ends A and B (5 inch x 3 inch)were in contact to the blood feeding chamber to allow for mosquito landing and probing. The Hemotek membrane feeding system was used to feed the blood sucking mosquitoes through an artificial membrane. KEMRI breeds and rears the *Anopheles* mosquitoes for malaria research. Animal blood (cow blood) used in the study as a meal attractant (Source: the blood stock in Kenya Medical Research Institute (KEMRI) which was used as a regular meal for mosquitoes). The animal blood (cow) was maintained at 38.60°C, the body temperature of a healthy cow (Kou *et al.,* 2017). The blood chamber which is an aluminium cylindrical container was loaded with fresh blood by means of a pipette through the ports at its back. The ports are covered with a removable rubber material. The mosquitoes pick up cues that indicate presence of animals or humans through vision for spotting the host and thermal sensory information to detect body heat(McMeniman *et al.,* 2014; EarthSky in Earth, 2015). The loaded blood in the chamber was covered with an artificial membrane (cellulose membrane with pores) and connected to the cage netting as given in Figure 2 and 3.

**Figure 2:**
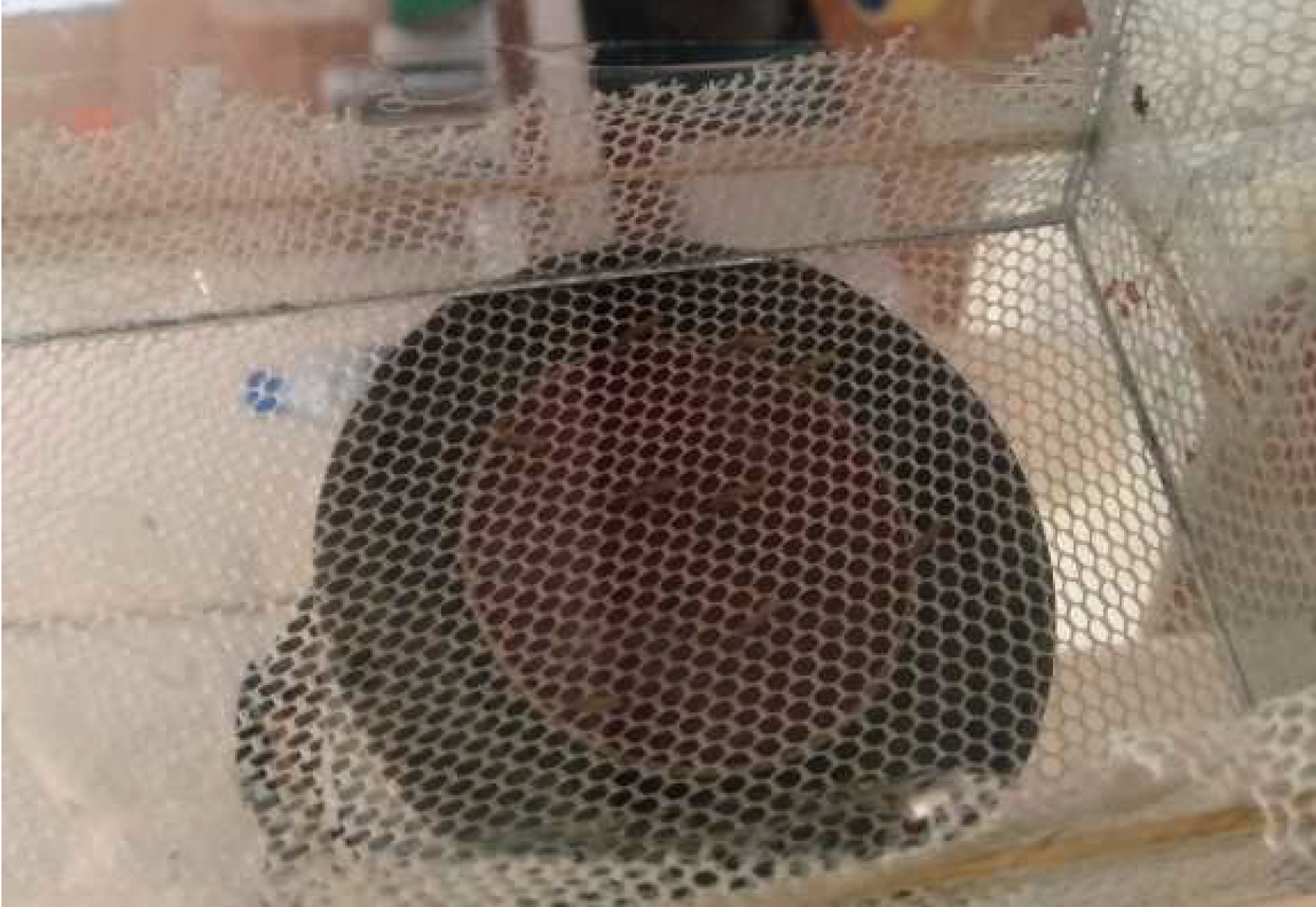
Mounted blood chamber on the net by means of retort stand

**Figure 3:**
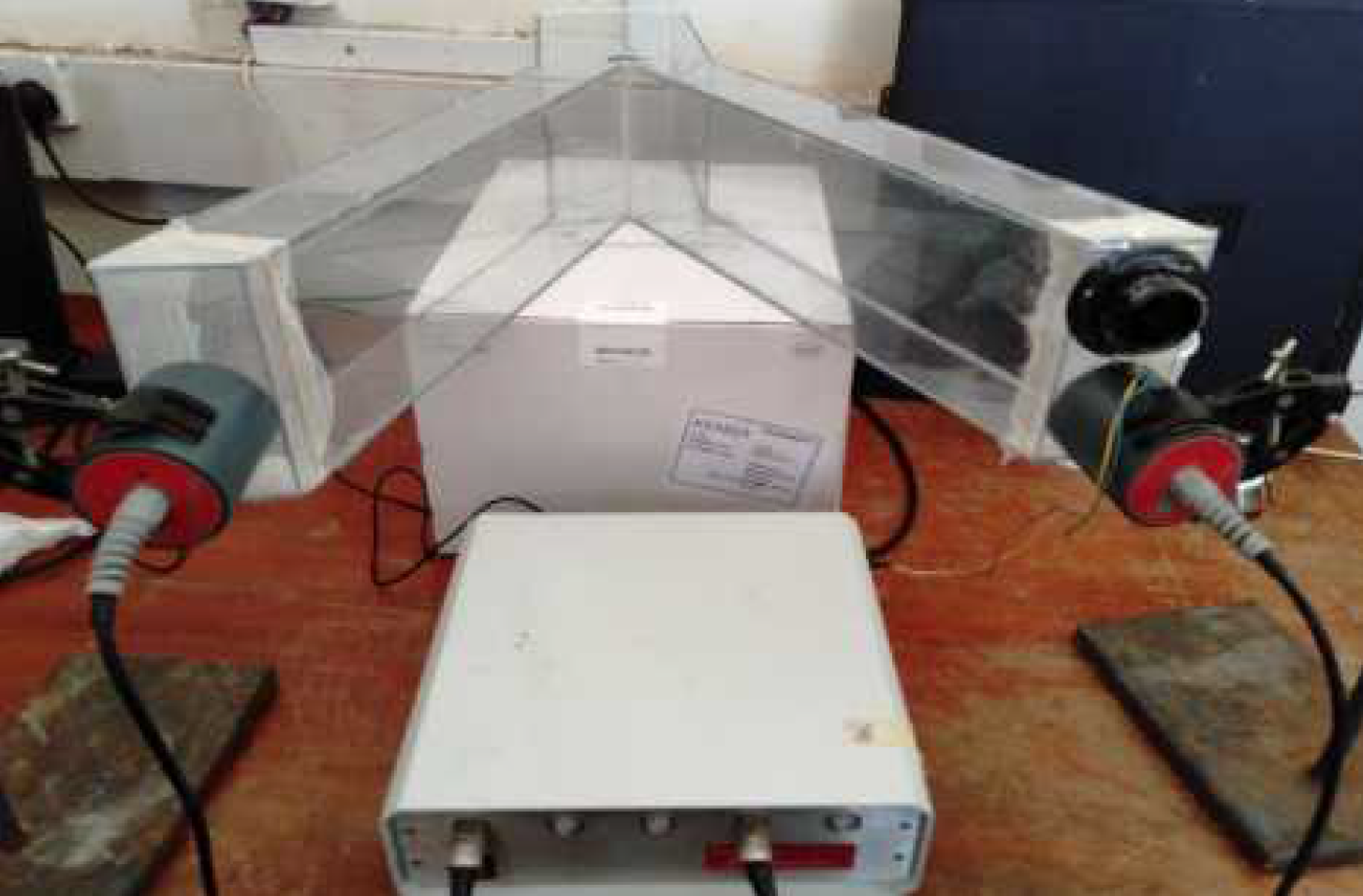
Bioassay Set-up with Cage A as Treatment cage

The animal sound (treatment) was allowed through the netting on cage A and B interchangeably as given in Figure 3.

Fifty laboratory reared mated female *A. gambiae* were allowed to the neutral chamber by means of an aspirator through the opening on the top of the cage in Figure 3. The animal sounds in the range of 10-34 kHz, 35-60 kHz and 61-90 kHz were played separately at a duration of 1,200 s, and observations of the mosquito behaviour made and recorded at an interval of 120 s. The startle responses exhibited by mosquitoes on exposure to the attractant and sound were typically based on the number of mosquitoes that approached the attractant; landed; landed and probed; or landed, probed, and bit the repellent-treated blood meal (Barnard, 2005). The study was based on the in vitro method and the ASTM E951-94 repellent procedures (Buescher *et al.,* 1983) with the treatment being the various frequency bands of animal sounds. A set of 50 mated and unfed female *A. gambiae* samples, 3 - 5 days old were transferred into the glass cage by means of an aspirator through a hole perforated on the top surface of the glass cage. The number of female *A. gambiae* that approached the blood meal; landed; landed and probed were determined in a time interval of 120 s were observed and recorded. Duration was determined using a timer. The data obtained was used to determine the protection index of the sound of *O. tormota, male A. gambiae* and *D. leucas.* The cage with no sound was the control for the bioassay study as given in Figure 4. The female *A. gambiae* startle response results determined from the bioassay study involving the sound of the sound of *male A. gambiae* and *D. leucas* were statistically compared with the results determined and reported for the sound of *O. tormota.*

**Figure 4:**
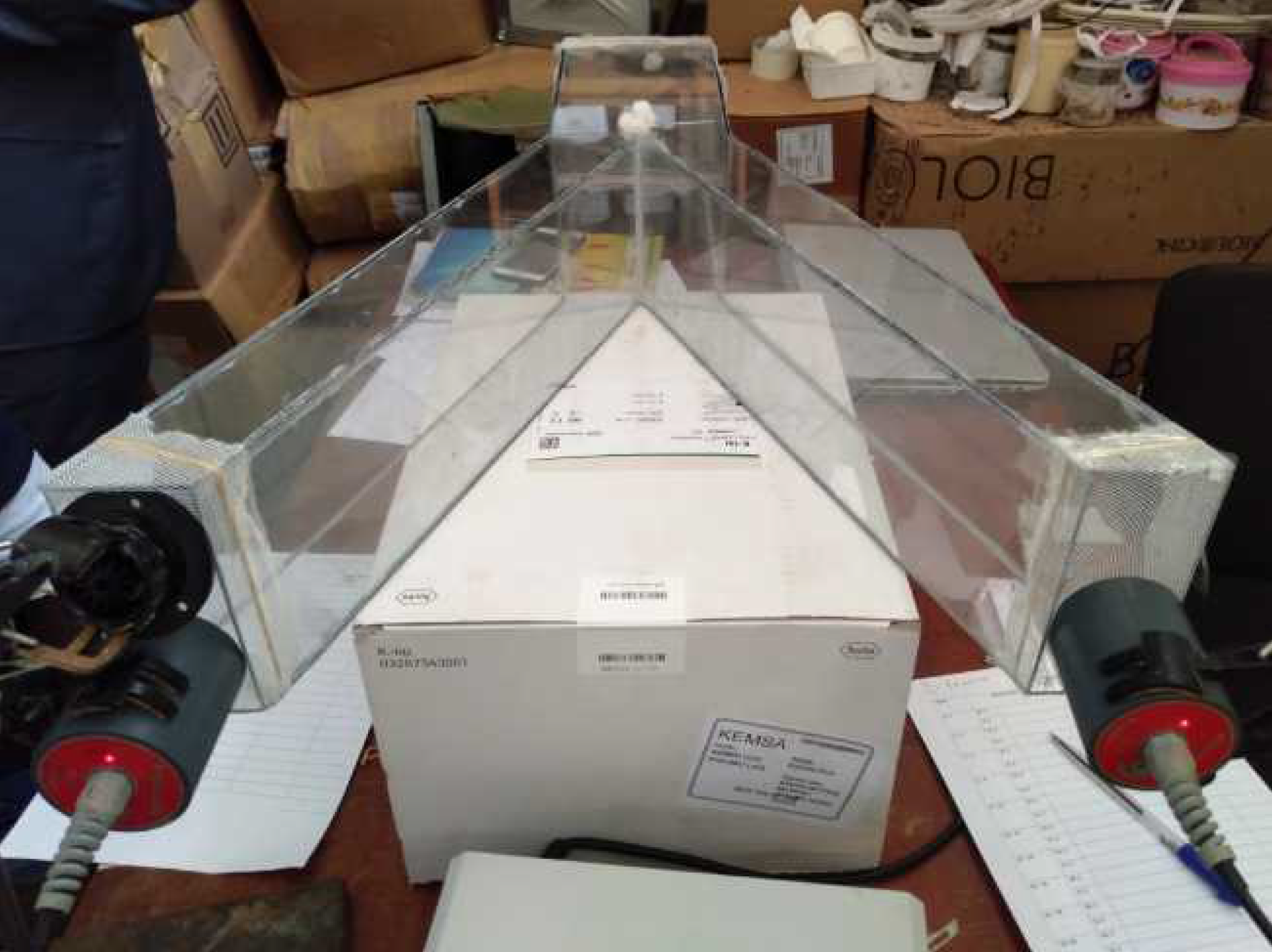
The Bioassay set up B as Treatment cage

## 3. Results and Discussion

### 3.1. Investigation of the Startle Response of the African female *Anopheles gambiae* evoked by Natural Sound of male mosquito, *Anopheles gambiae*, *Odorrana tormota* and *Delphinapterus leucas*

#### 3.1.1. Determination and analysis of the landing rates and behavioural startle responses of the mated female *A. gambiae* on food attractant evoked by the 10-34 kHz individual sound of the male mosquito, *A. gambiae*, *O. tormota* and *D. leucas*

The 10-34 kHz filtered sounds of *O. tormota, male A. gambiae* and *D. leucas* were used in the study informed by the initial startle effect of the sound of *O. tormota* in past studies (Mang’are *et al*., 2012). The recent studies on startle effect yielded a startle response of 34.12 % but limited to initial behavioural response without a mosquito attractant and was not based on protection index (PI). The study with the current bioassay setting involved new sound clips of *O. tormota* which required further investigation of to determine the protection index. A total of 50 female *A. gambiae* were allowed into the neutral cage by means of an aspirator at a time and allowed to enter the open Control bioassay chamber or the treatment bioassay chamber with equal chance of probability as shown in Figure 3 and 4.

##### (i). Determination and analysis of the landing rates and behavioural startle responses of the mated female *A. gambiae* on food attractant evoked by the sound of *O. tormota*

The control bioassay experiment did not involve any sound and was conducted in one chamber (control cage). In the control cage, the female *A. gambiae* were observed flying freely towards the blood meal. The female *A. gambiae* flew about and rested on the glass walls and net at normal posture, projecting its abdomen in air at about 45°. Low flights and landings on blood meal was observed. Fully fed whose abdomen was engorged and appeared red flew lowly and rested at the base and at times on the wall with minimal movements. Some fully fed mosquitoes flew out of the control chamber and rested on the net in the neutral cage as given in Figure 5. The low flights were attributed to the weight of the mosquito which had increased. Free flight from the neutral cage to control cage and back among the mosquitoes was noted. The mosquitoes appeared relaxed. However, the bioassay cage in which the warm blood meal was treated with the 10-34 kHz sound of *O. tormota* showed a reduced number in the mosquitoes approaching and/or landing on the blood meal. Restlessness and rubbing of wings with hind legs was observed. The mosquitoes which had entered the treatment cage exhibited flights along the walls, bouncing on the wall and moved towards the neutral chamber. The mosquitoes which managed to land, probe and feed flew out of the treatment cage to the neutral cage resting on the net. Two mosquitoes were observed shaking the body while feeding while others landed on the meal and flew away without feeding. The frequency of flights in and out of the chamber increased with the fully fed mosquitoes exhibiting immobilised. This was attributed to neural stress and fear of predation.

**Figure 5:**
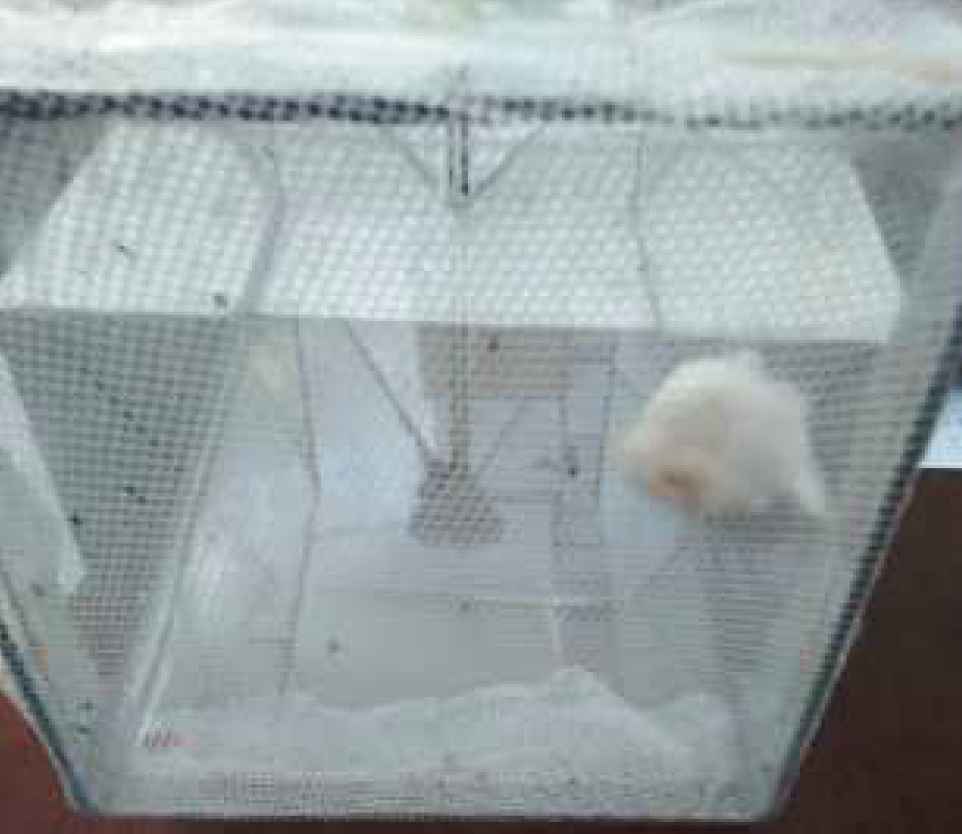
Mosquitoes resting on the net in the neutral chamber

All the mosquitoes in the control and treated chamber including those that landed, successfully or unsuccessfully probed and fed were considered to have approached the blood meal. Based on this premise, the number of the female *A. gambiae* that approached the control chamber exceeded the number of the mosquitoes that approached the treatment cage significantly (**p = 8.5381 x 10^−6^ <<< 0.05**) as shown in Figure 6 and correlated positively low (**r = 0.3715**). The sound of *O. tormota* yielded a Protection index (PI) of **26.76** % based on the number of mosquitoes that approached the blood meal, which was 7.36 % below the startle response of 34.12 % for the sound of *O. tormota* in the same frequency range. The difference in repellency determined through One-Sample T-test Statistics was highly significant (p = 4.4992 x 10^−12^ <<<< 0.05).

**Figure 6:**
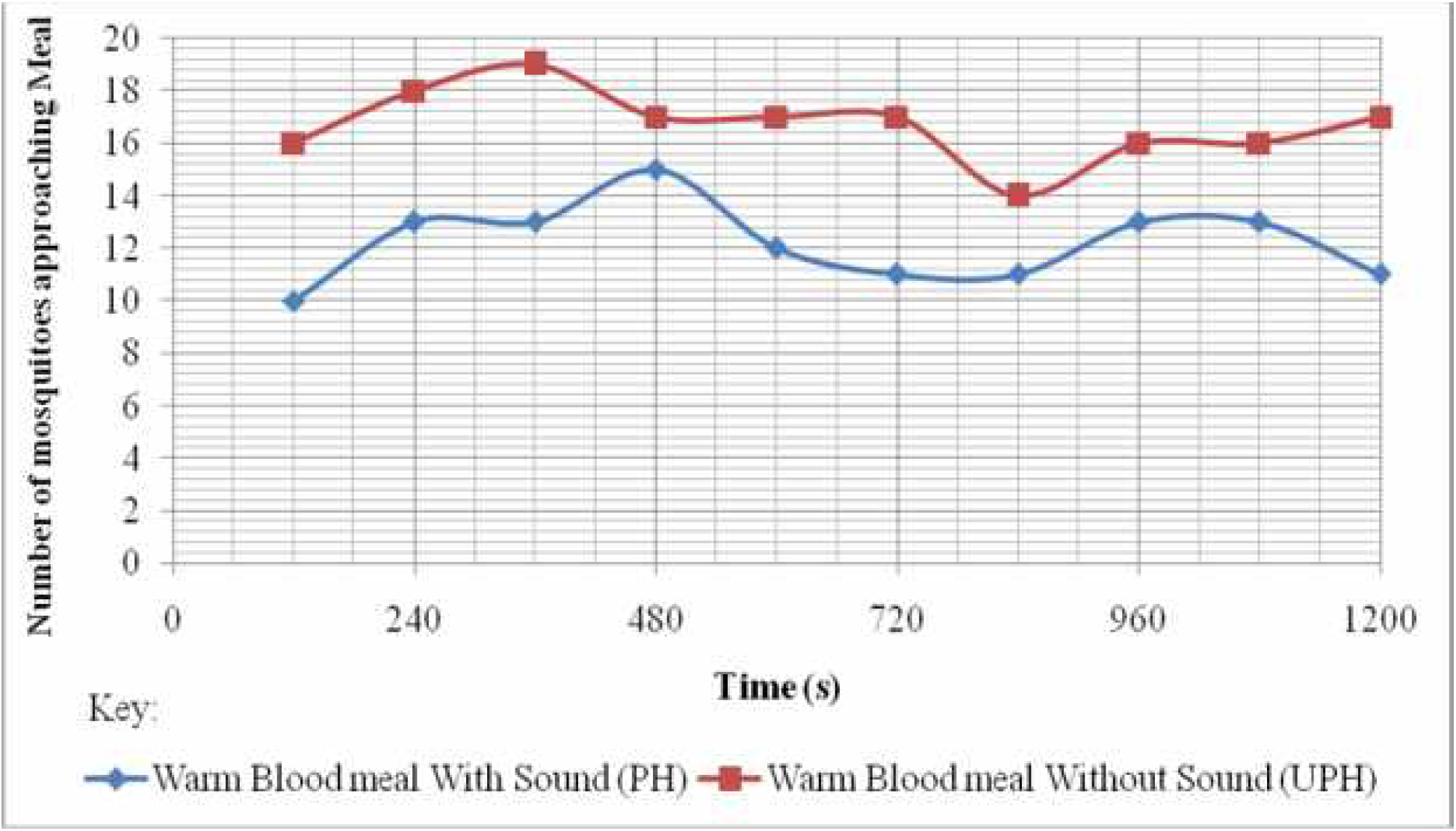
Number of mosquitoes approaching the control and treatment cage for the sound of *O. tormota*

The number of mosquitoes that landed, probed and fed on the blood meal in the treatment cage was lower compared to the number of mosquitoes that landed, probed and fed on the blood meal in the control cage yielding a protection of **40.24** % as given in Figure 7. The paired sample T-test comparison of number of mosquitoes that landed, probed and fed on the blood meal in the treatment cage to ones that landed, probed and fed on the blood meal in the control cage showed a highly significance difference with significance value p = 1.9362 x 10^−5^ with a strong positive correlation (Pearson’s correlation value r = 0.9599). The protection index evoked by the sound of *O. tormota* was 6.12 % higher than the reported startle response in female *A. gambiae* in the same frequency band, a difference which was significantly high (p = 1.0225 x 10^−10^)as determined through One-Sample T-test Statistics.

**Figure 7:**
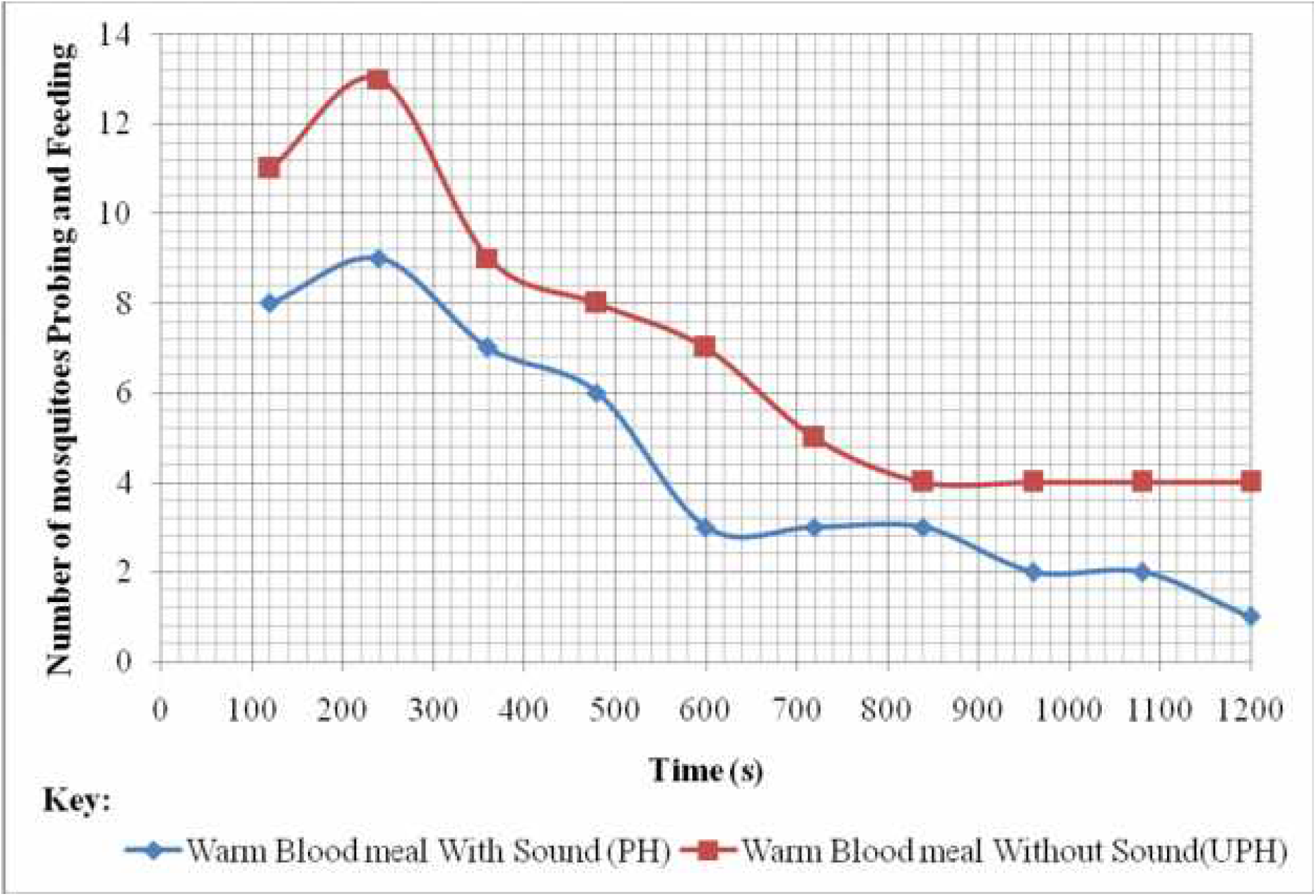
Number of mosquitoes feeding in the control and treatment cage for the sound of *O. tormota*

The protection index evoked by the sound of *O. tormota* significantly exceeded the repellency due to the sound from the Anti-Pic^®^ (electronic mosquito repellency) by 9.94 % which had been claimed to repel 30.3 % mosquitoes (Andrade and Bueno, 2001). Also, the protection index evoked by the sound of *O. tormota* was higher than the repellency determined from experiments with functioning electronic mosquito repellents (EMR) that mimicked calls from bats and male *A. gambiae* in by 20.24 % (Andrade and Bueno, 2001; Center for the Advancement of Health, 2007; Enayati *et al*, 2010). The 10-34 kHz frequency range of the sound of *O. tormota* was pulsate in nature with a maximum acoustic energy and power of 6.7437 Pa^2^s and 110.10 dB respectively. There were 2,654 calls each lasting for a mean duration of 0.0439 s with mean measurements of 14.02 kHz, 19.40 kHz and 95.68 Pa for peak frequency (maximum entire), maximum frequency (maximum entire) and peak amplitude (mean entire) respectively for the sound of *O. tormota*. The minimum and mean acoustic energy of the 10-34 kHz frequency range of the sound of *O. tormota* was 0.00025 Pa^2^s and 0.4442 Pa^2^s respectively. The highly pulsate sound declined sharply giving a power deviation, ΔP = 38.5 dB. The 2,654 calls recorded a maximum and mean call duration of 0.4198 s and 0.0439 s respectively. The mean and maximum measurements of the bandwidth (mean entire) was 5.29 kHz and 16.90 kHz respectively.

##### (ii). Determination and analysis of the landing rates and behavioural startle responses of the mated female *A. gambiae* on food attractant evoked by the sound of the male mosquito, *A. gambiae*

The control bioassay chamber and treatment bioassay chamber were connected to the feeding membrane through the netting. The food attractant on the treatment bioassay cage was exposed to the 10-34 kHz recorded and filtered sound of the male mosquito, *A. gambiae.* The female *A. gambiae* remained rested in the neutral chamber before deciding to move to the control or treatment bioassay chamber. The female *A. gambiae* were observed flying freely in and out of the control chamber. Minimum mosquito movement was observed in the control bioassay chamber. A total of six unfed female *A. gambiae* rested on the wall and the floor of the chamber. The female *A. gambiae* rested by projecting their abdomen in air at about 45°. Fully fed mosquitoes rested on the wall and the floor of the control bioassay chamber. Mosquitoes in both control bioassay chamber and treatment bioassay chamber rubbed their wings with their limbs to clean them. In the treatment bioassay chamber, the female *A. gambiae* were observed flying about, with one appearing disturbed. Two mosquitoes settled at the edge between the neutral chamber and treated chamber after failing to land, probe and feed the blood meal. One mosquito flew from the treatment bioassay cage to the control cage. Some mosquitoes were observed not to be fully fed. The number of mosquitoes inside the chamber, either the Control or treated chamber, were considered to have approached the blood meal. A comparison of the number of the female *A. gambiae* approaching meal in the control bioassay cage exceeded the number of the female *A. gambiae* in the treatment bioassay cage except at the 600^th^ second where the number of *A. gambiae* mosquitoes in both chambers were equal as shown in Figure 8. The equality in the number of the female *A. gambiae* approaching the blood meal in the control bioassay cage and the treatment was attributed to the pulsate nature of the sound and weak acoustic energy and power during the 600^th^ second.

**Figure 8:**
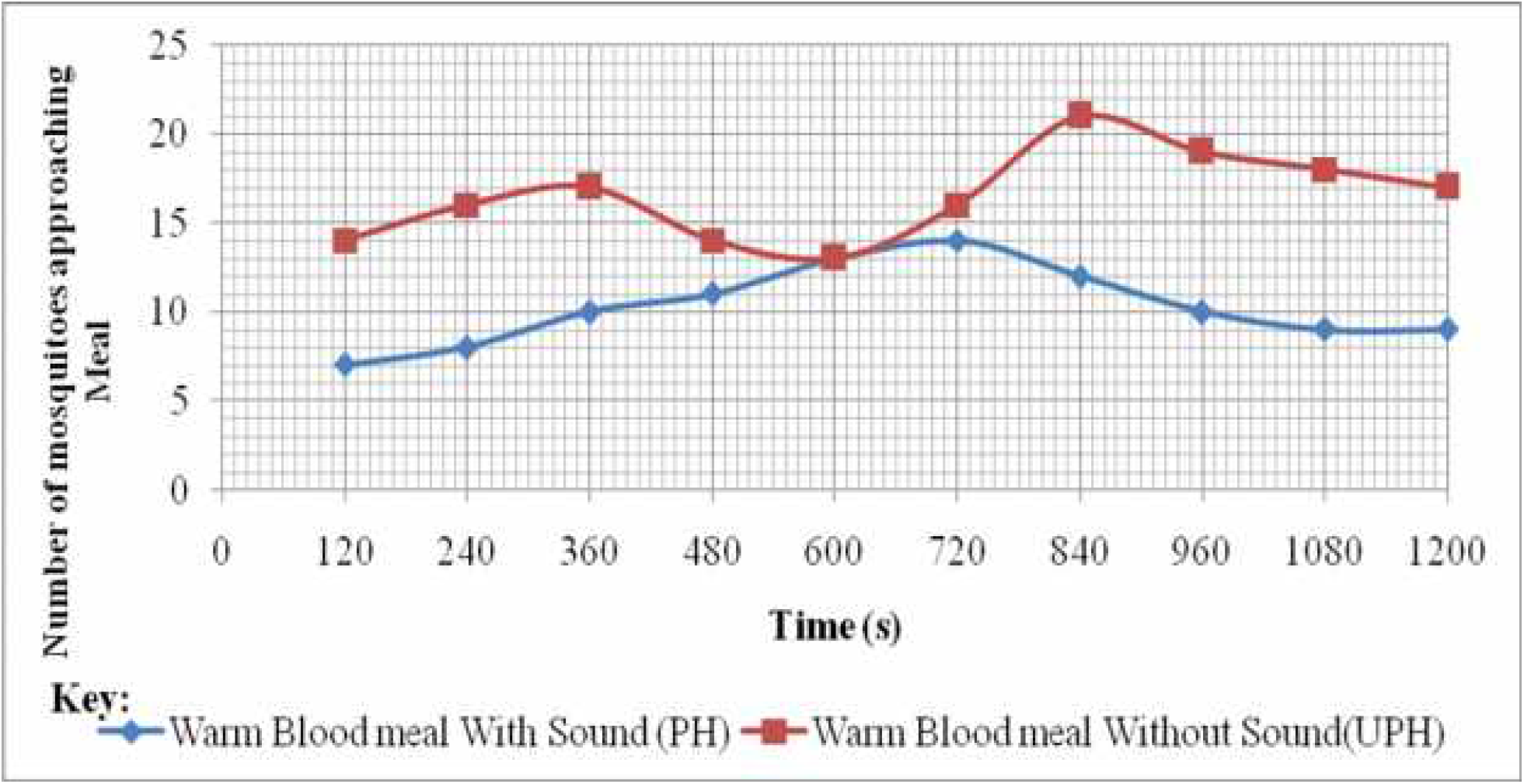
Comparison of the number of mosquitoes approaching the control and treatment cage

The reduction in the number of female *A. gambiae* approaching the warm blood meal which was treated with the 10-34 kHz sound of male mosquito, *A. gambiae* was highly significant (p = 2.1436 x 10^−4^ <<< 0.05) and correlated positively low (r = 0.0102). The protection index due to the 10-34 kHz sound of male mosquito, *A. gambiae* was **36.24 %** exceeded the repellency owing to the 10-34 kHz sound of *O. tormota* by **9.48 %,** with both studies based on the number of mosquitoes that approached the respective blood meals. However, difference in the protection index due to the sound of *A. gambiae* and protection index due to the sound of *O. tormota* was not significant (p = 0.1740 > 0.05) and correlated negatively low (r = −0.0235). The protection index due to the sound of the male *A. gambiae* was 2.12 % above the reported startle response of **34.12 %** of to the sound of *O. tormota* in the same frequency band, a difference that was not significant (p = 0.7209 > 0.05). Also, the protection index due to the sound of the male *A. gambiae* exceeded the repellency due to the sound from the Anti-Pic^®^ (electronic mosquito repellency) by 5.94 % though not significantly (p = 0.3285 > 0.05) as determined through One-Sample T-test Statistics.

The number of the mated female *A. gambiae* that landed, probed and fed on the blood meal in the treatment cage was lower compared to the number of mosquitoes that landed, probed and fed on the blood meal in the control cage yielding a protection index of **42.73 %** as given in Figure 9.

**Figure 9:**
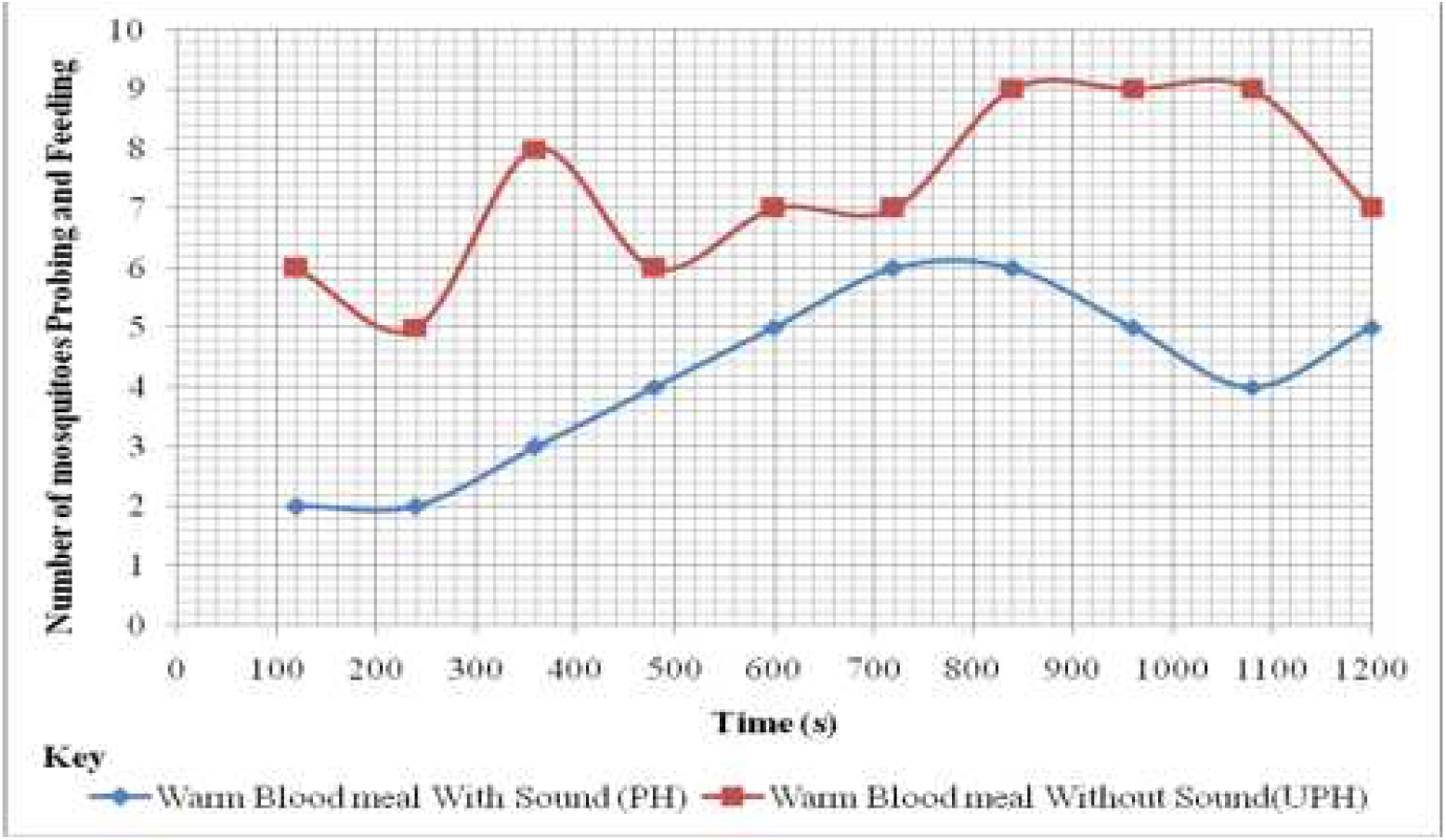
Number of mosquitoes feeding in the control and treatment cage for the sound of the male *A. gambiae*

The paired sample T-test comparison of number of mosquitoes that landed, probed and fed on the blood meal in the treatment cage to the number of mosquitoes that landed, probed and fed on the blood meal in the control cage indicated a highly significance difference in repellency (p = 5.3440 x 10^−5^ < 0.05) with a strong positive correlation (Pearson’s correlation value r = 0.5522). The sound of *O. tormota* yielded protection index which was **2.49 %** lower compared to the sound of the male *A. gambiae* for experiments based on the number of mosquitoes that landed, probed and fed on the blood meal in both control and treatment bioassay cage. The difference in repellency elicited by the sound of the male *A. gambiae* compared and the sound of *O. tormota* was not significant (0.1740 > 0.05). However, the difference in repellency due to the sound of the male *A. gambiae* was 12.43 % higher than the claimed repellency due to the sound from the Anti-Pic^®^ electronic mosquito repellent and the difference was significantly high (p = 5.3440 x 10^−5^) as determined through One-Sample T-test Statistics. The sound of the male *A. gambiae* in the 10-34 kHz frequency range was minimally pulsate in nature with a maximum acoustic energy and power of 0.9173 Pa^2^s and 84.40 dB respectively, giving a marginal power deviation, ΔP = 9.40 dB. The maximum and mean of the bandwidth (mean entire) was 28.30 kHz and 24.61 kHz. The minimum acoustic energy of the sound of *O. tormota* exceeded the sound of the male *A. gambiae* by 0.00011 Pa^2^s. However, the maximum and mean acoustic energy of the sound of *O. tormota* exceeded the sound of the male *A. gambiae* by 5.8264 Pa^2^s and 0.4366 Pa^2^s respectively. The difference in acoustic energy of the sound of male mosquito, *A. gambiae* and *O. tormota* was highly significant with significant values, p = 4.4047 x 10^−50^ < 0.05 and the parameters correlated positively low (r = 0.0119). The sound of the male *A. gambiae* was composed of 41,135 calls whose maximum and mean duration was less than that of the sound of *O. tormota* by 0.1388 s and 0.0402 s respectively. The acoustic power of the sound of the *O. tormota* declined drastically compared to that of the sound of the male *A. gambiae* giving a giving a power deviation difference of 29.10 dB. The minimal power difference, pulsate natured signal, higher minimum acoustic energy, many short duration calls and wide bandwidth in the sound of the male *A. gambiae* evoked greater repellency compared to the sound of the *O. tormota.*

##### (iii). Determination and analysis of the landing rates and behavioural startle responses of the mated female *A. gambiae* on food attractant evoked by the sound of *Delphinapterus leucas*

The 10-34 kHz frequency band was filtered from the entire spectrum of the sound of *D. leucas* and used for the bioassay study. In the control bioassay chamber, the blood meal was not treated with sound. The mosquitoes flew about, to and from the neutral cage with some unfed mosquitoes resting normally on the walls of the cage during the first 120 s. Ten mosquitoes were observed feeding with no signs of disturbance. There was minimum movement by the mosquitoes in the cage after 240s. Two fully fed mosquitoes were observed resting on the cage after 480 s. Notably, one fully fed mosquito flew from the treated chamber to the control chamber then settled in the neutral cage during the 600^th^ second. Minimum movement and activity with one unfed mosquito rubbing its wings with hind legs was observed in the chamber up to 1080^th^ second. During the last 120s, two fully fed mosquitoes were seen resting in the cage.

In the treatment cage, the mosquitoes were seen approaching the meal and landing and flying away along the wall, with others flying around the meal without successful landing during the first 120 s. Between 120s and 240s, minimal movement was observed with on unfed mosquito flying into the chamber and resting on the wall of the cage. During the 480-600s duration, two fully fed mosquitoes flew from the treatment bioassay cage to the neutral cage where it rested on the net. One unfed mosquito flew from the neutral chamber into the treated chamber but flew back to the neutral cage immediately. Other unfed mosquitoes rested to the walls and floor with minimum movement during the same duration. Bouncing along the net and on glass walls associated with knocks was observed during the 840-1200s duration.

The number of the female *A. gambiae* approaching the blood meal in the control bioassay cage were more than the number of the female *A. gambiae* in the treatment bioassay cage except during the 240^th^ second where the number of mosquitoes in the treatment cage exceeded the control as given in Figure 10. Instances of attraction were confirmed during the 240^th^ second giving an instantaneous protection index of −6.67 % due low acoustic energy and power. The sound was least pulsate in nature.

**Figure 10:**
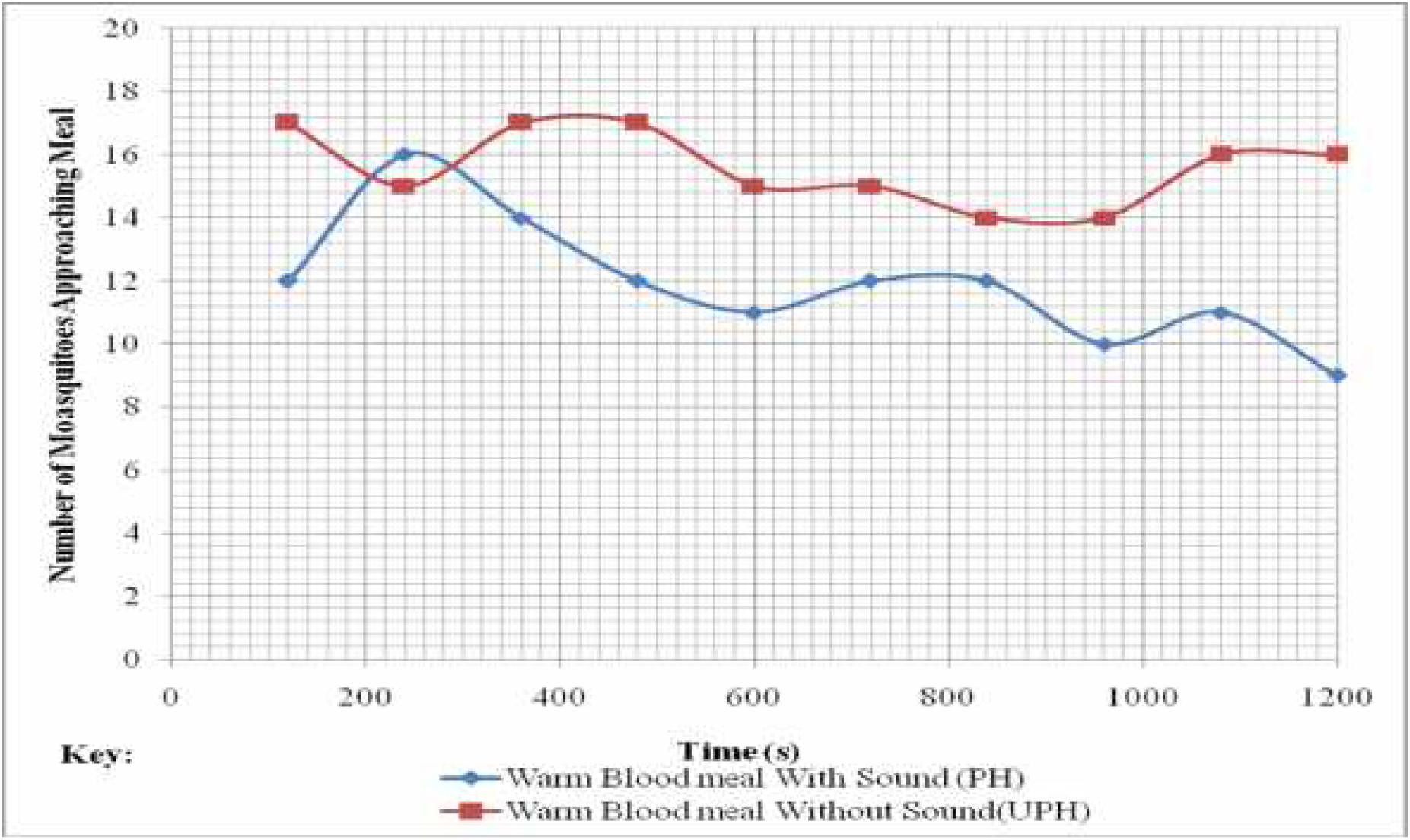
Number of mosquitoes approaching in the control and treatment cage for the sound of *D. leucas*

The difference in the number of mosquitoes approaching the meal in the control bioassay cage and the treatment bioassay cage was highly significant (p = 4.2749 x 10^−4^ < 0.05) and the parameters correlated positively low (r = 0.1250). The overall protection index based on the number of mosquitoes approaching the blood meal in the control and treatment bioassay cages was **23.43 %** which was less than the protection index for sound of male mosquito, *Anopheles gambiae* and *Odorrana tormota* by **12.81 %** and **3.33 %** respectively. The number of mosquitoes that landed, probed and fed on the blood meal in the treatment cage were less than the number of mosquitoes that landed, probed and fed on the blood meal in the control cage except during the 240^th^, 840^th^ and 1080^th^ seconds where they were equal in number as given in Figure 11. The number of mosquitoes that landed, probed and fed on the blood meal in the treatment cage during the 960^th^ second exceeded the number of mosquitoes that landed, probed and fed on the blood meal in the control cage.

**Figure 11:**
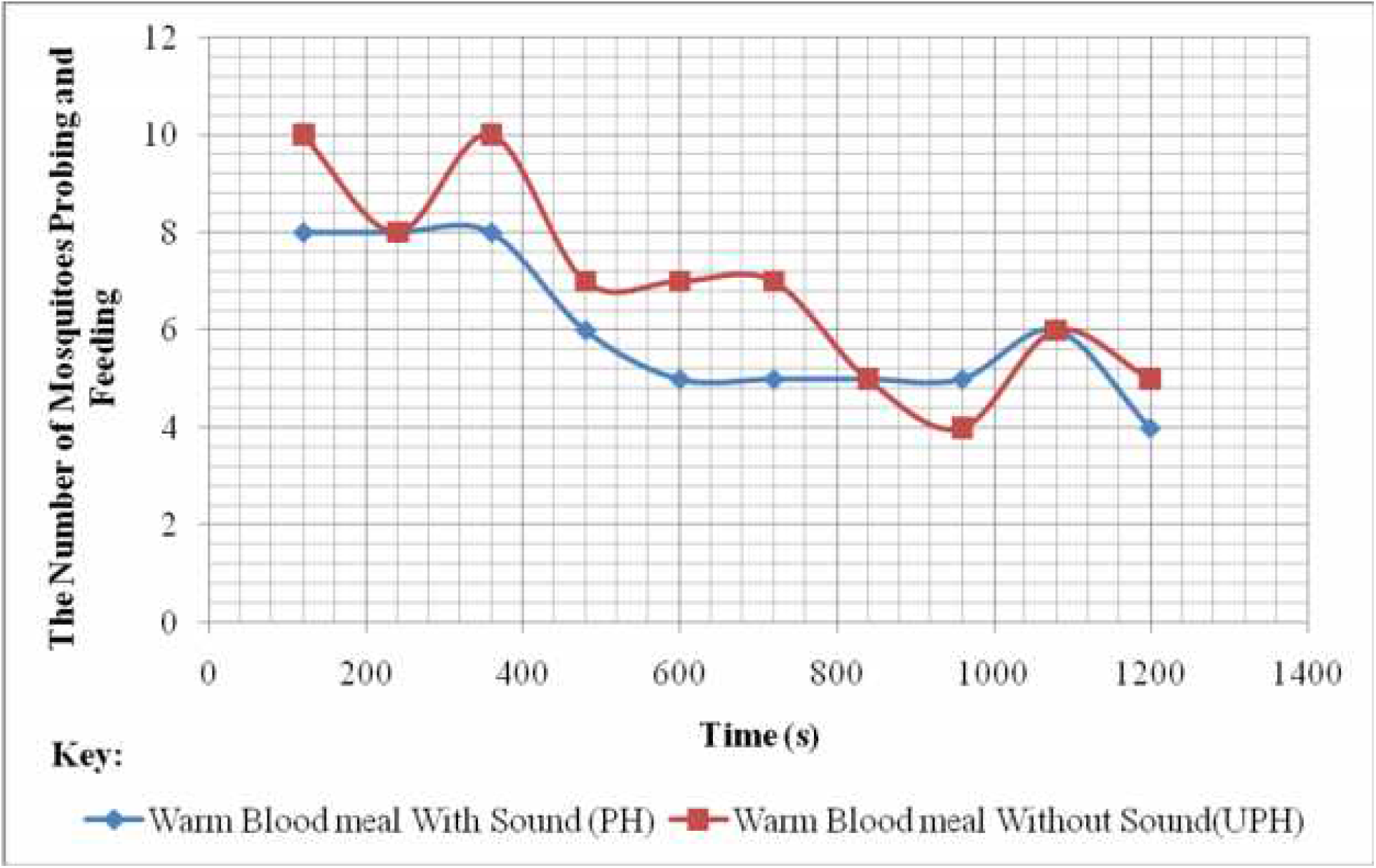
Number of mosquitoes Feeding in the Control and Treatment cage for the sound of *D. leucas*

The protection index based on the number of mosquitoes that landed, probed and fed on the blood meal in the treatment cage and the number of mosquitoes that landed, probed and fed on the blood meal in the control cage for the sound of *D. leucas* was **10.64 %**, lowest by **32.09 %** and **29.60 %** from the protection index due to the sound of male mosquito, *A. gambiae* and *O. tormota* respectively. The paired sample T-test comparison of number of mosquitoes that landed, probed and fed on the blood meal in the treatment cage to the number of mosquitoes that landed, probed and fed on the blood meal in the control cage for the 10-34 kHz sound of *D. leucas* showed a highly significance difference in repellency (p = 0.02940 < 0.05) with a strong positive correlation (Pearson’s correlation value r = 0.8466). The recorded sound of *D. leucas* yielded repellency which was **19.66 %** lower than the claimed **30.3 %** repellency due to Anti-Pic^®^ which is an electronic mosquito repellent (Andrade and Bueno, 2001). A one sample T-test comparison of the difference in the protection index due to *D. leucas* with the claimed **30.3 %** repellency due to Anti-Pic^®^ and **34.12 %** startle response due to the sound of *O. tormota* was highly significant with significant values, p = 0.0048 and 0.0016 respectively. The 10-34 kHz sound of *D. leucas* recorded the least minimum acoustic energy of 0.0001 Pa^2^s compared to the sound of the male *A. gambiae* and *O. tormota*. The maximum acoustic energy of the sound of *D. leucas* exceeded the acoustic energy of the sound of male *A. gambiae* by 1.37011 Pa^2^s though less than the acoustic energy of the sound of *O. tormota* by 4.45637 Pa^2^s. The maximum and mean bandwidth (mean entire) in the 10-34 kHz sound of *D. leucas* was 25.30 kHz and 22.11 kHz respectively which exceeded the corresponding parameters of the sound of *O. tormota* but less than the sound of the male *A. gambiae*. The minimum and maximum acoustic power of the sound of *D. leucas* was 50.00 dB and 71.00 dB respectively, giving a power deviation difference of 21.00 dB. The call duration of the 15,299 calls lasted for a mean duration of 0.0013 s. The 10-34 kHz sound of the male *A. gambiae* provided the highest protection index compared to other sounds under study as shown in Figure 12.

**Figure 12:**
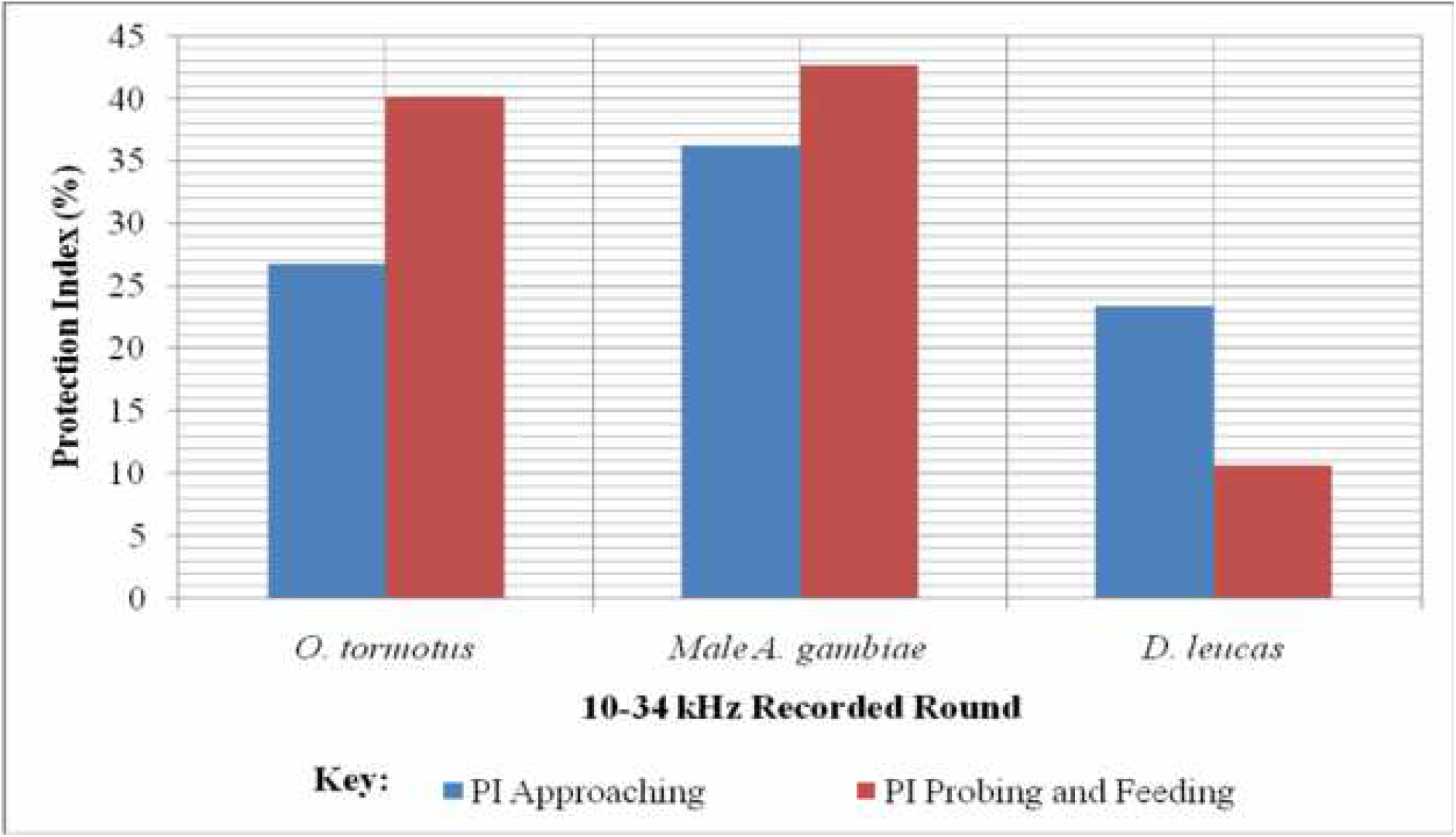
Comparison of the protection indices (PI) of animal sounds

The pulsate and almost steady trend of the acoustic power in the sound of the male *A. gambiae* and wide mean of the bandwidth (mean entire) provided superior parameters for the highest protection index and least landing rates as shown in Figure 12 and 13. The 10-34 kHz frequency range for the sounds of the *A. gambiae, O. tormota* and *D. leucas* elicited 2.10 landings /minute, 2.20 landings/minute and 3.00 landings/minute respectively. Also the deviation in power was minimal within the same frequency range. The ultrasonic components caused neural stress and a refractory behaviour to the male sound. However, the protection index due to the sound of the male *A. gambiae* was higher than the reported electronic mosquito repellent devices and that of *O. tormota* earlier investigated. The methodology employed in establishing repellency in female *A. gambiae* due to the sound of *O. tormota* in earlier studies differed with the current approach which is an improved methodology. The sound of *D. leucas* evoked the least repellency due to the minimal pulsate nature of the sound, low maximum and mean acoustic energy and low acoustic power. The sound of *O. tormota* was highly pulsate in nature with the greatest maximum acoustic energy and power. However, the steep slope in acoustic power, narrow bandwidth and wide power deviation, ΔP = 38.5 dB lowered the protection index of the sound of *O. tormota* in the 10-34 kHz frequency range.

**Figure 13:**
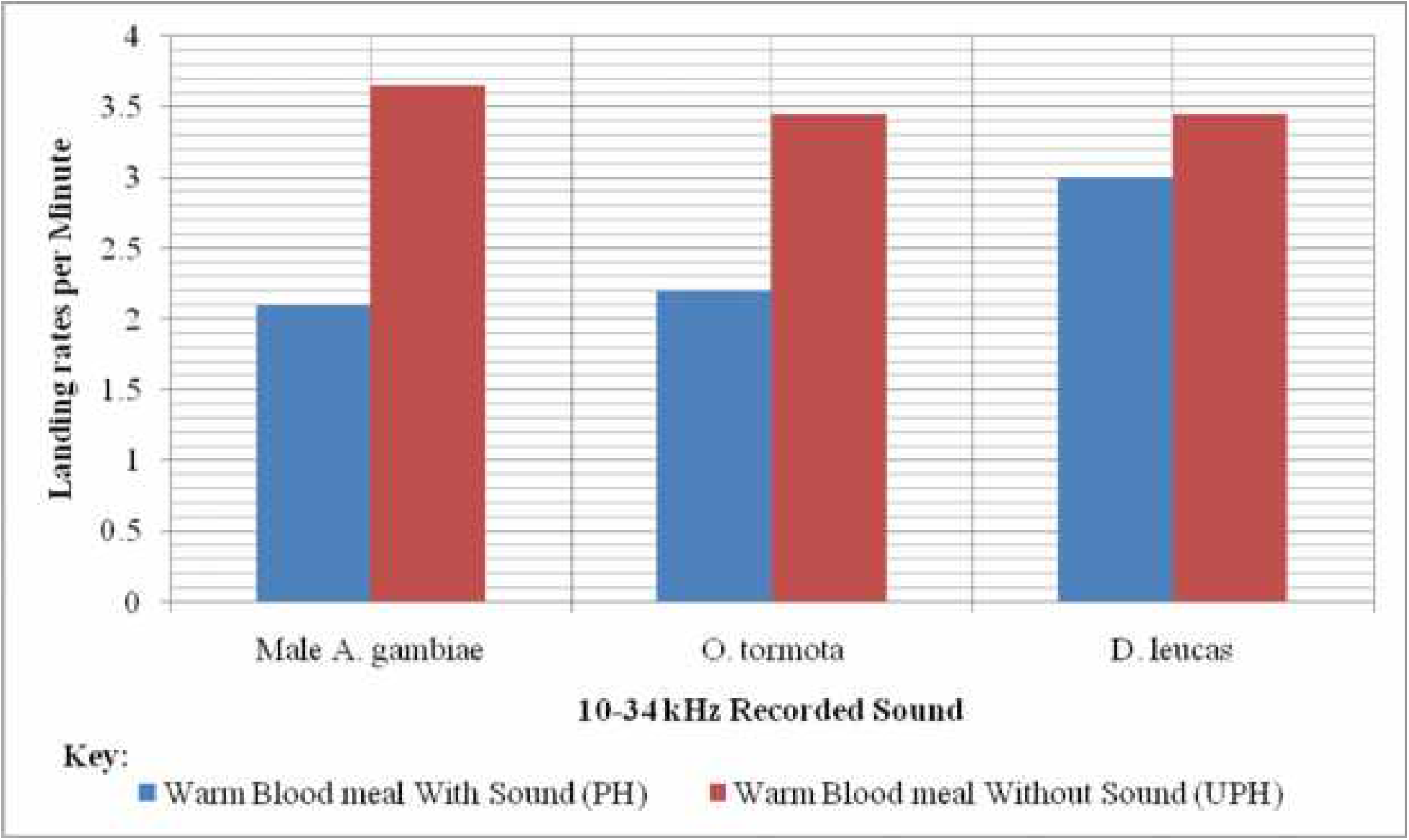
Comparison of the Landing rate evoked by Animal sounds

## Conclusion

The sounds of the male *A. gambiae, O. tormota* and *D. leucas* evoked different degrees of protection against the female *A. gambiae* with the sound of the male *A. gambiae* yielding the greatest protection index of **42.73 %** with the least landing rate of **2.10** landings/minute. The high protection index was attributed to the pulsate and almost steady acoustic power, wide mean bandwidth (mean entire) and minimal deviation between the maximum and minimum acoustic power within the 10-34 kHz frequency range.

## Acknowledgement

I am greatly indebted to Egerton University and Masinde Muliro University of Science and Technology for giving me an opportunity to conduct the research. Many thanks are extended to Prof. Feng, Raimund Specht of Avisoft Bioacoustics, Pettersson Elektronik AB, Cornell Lab of Ornithology, Bernard Agwanda of the National Museums (Kenya) and Prof. Herve Glotin of Institut Universitaire de France for their kind donations and technical support. I also thank the Director and the staff of KEMRI (Kisumu) for allowing us to use their facilities compounded with their technical input.

